# Spatial relationship of urban carbon neutrality in Yangtze River Basin

**DOI:** 10.1101/2025.03.14.643198

**Authors:** Qing Li, Xiaohe Meng, Xiangyan Chen, Yu Liu

## Abstract

Carbon neutrality needs to be implemented spatially in order to be realized concretely. The distribution of carbon sources and sinks across the national territory is closely related to many factors, such as natural geographical resources, levels of economic and social development, and cultural and historical heritage. It is difficult for many regions to achieve carbon neutrality through their own efforts alone. This article believes that only by achieving local optimal carbon neutrality through a reasonable spatial organization can emissions be ultimately reduced to zero on a national scale. Under the connection of rivers, the basin forms an ecosystem and an economic system with independent characteristics. Carbon sources and sinks within the basin exist within the ecological and economic space of the basin, forming a unique carbon neutrality spatial pattern. This study starts with the spatial distribution status, features and relationships of carbon neutrality elements in the cities of the Yangtze River Basin, and divides the cities into eight types based on their development level, emission level and forest resources, such as high-emission and low-carbon sink developed cities and low-emission and high-carbon sink underdeveloped cities. Different types of cities are proposed with carbon neutrality paths and spatial organization schemes, and it is suggested that the spatial organization demonstration of carbon neutrality in the cities of the Yangtze River Basin can provide a reference for carbon neutrality on a national scale.

## 1. Introduction

To address climate change, the Paris Agreement proposes achieving net-zero carbon emissions by 2050 and keeping the global surface temperature rise within 2°C above pre-industrial levels by the end of this century. Currently, China’s greenhouse gas emissions account for about 30% of the global total, and although it is the world’s second-largest economy, it still faces enormous development pressures, with per capita income below the world average and only one-sixth that of the United States. Additionally, 600 million people have a disposable income of only 1,000 RMB. China’s industrial structure is heavily reliant on heavy industry, and its energy structure is dominated by high-carbon fossil fuels. Compared to developed countries, the uneven development of its economy and society makes it much more challenging for China to achieve carbon peaking and carbon neutrality while maintaining economic growth (Fowler et al., 2020). However, President Xi Jinping has pledged to increase China’s nationally determined contributions with more powerful policies and measures, aiming to peak carbon dioxide emissions by 2030 and achieve carbon neutrality before 2060, setting clear targets for China to address climate change (Huang, 2020).

Achieving carbon neutrality involves both reducing carbon source emissions and increasing carbon sink absorption (Fowler et al., 2020). In the past, reducing emissions was the main approach to mitigate climate change. However, as the marginal cost of emissions reduction increases and the marginal benefit decreases, the difficulty in reducing emissions has grown (Tian et al., 2021). As a result, increasing carbon sink absorption has become increasingly important, especially after carbon peaking is achieved in 2030 (Lu, 2013; Yang, 2013). From the perspective of reducing carbon source emissions, reducing greenhouse gas emissions is “subtractive” by improving energy efficiency, reducing energy consumption, and increasing clean energy (Ping et al., 2020; Yang and Li., 2020). These measures require significant economic and societal adjustments, at great development costs

(Luo et al., 2020). After years of energy conservation and emission reduction efforts, China’s carbon intensity had decreased by approximately 48.1% from 2005 by the end of 2019, and non-fossil energy accounted for 15.3% of primary energy consumption (Tian et al., 2021; Xu et al., 2023). This has made a significant contribution to global emissions reduction (Sheng and Lu, 2016). From the perspective of increasing carbon sink absorption, increasing carbon sinks involves fixing carbon dioxide within plants or soil, thus reducing greenhouse gas concentrations in the atmosphere. This approach adds carbon sinks as contributions to carbon neutrality (Lu, 2013). Since the Kyoto Protocol, international climate policy has identified enhancing forest carbon sinks as a major way to achieve carbon neutrality (Zhang et al., 2021). The Paris Agreement in 2015 confirmed that enhancing carbon sinks and reducing emissions are effective ways to mitigate global warming. Starting from the 1980s, China has implemented the world’s largest afforestation project in arid and semi-arid areas in the north to alleviate desertification and control sandstorms. From 1977 to 2018, China’s total forest carbon storage increased from 4.050PgC (1Pg = 1015g) to 8.362PgC, an increase of 106.46%. Forest stand carbon storage increased from 3.938PgC to 7.669PgC, an increase of 94.74%. Forest stand carbon density increased from 35.24 Mg/ha (1Mg = 106g) to 42.63 Mg/ha^1^, making significant contributions to global efforts to address climate change.

Carbon neutrality must be implemented spatially to be realized concretely. In terms of the entire national territory, the distribution of carbon sources and sinks is closely related to factors such as natural geographic resources, economic and social development levels, cultural and historical inheritances, etc. Many regions cannot achieve carbon neutrality alone, and only rational spatial arrangements can realize optimal carbon neutrality across the country (Masahisa et al., 2001). Carbon neutrality can be achieved across the entire national territory or within specific regions, river basins, or even individual projects (Zhang et al., 2021). To achieve carbon neutrality targets, spatial organization and arrangements for carbon neutrality must be made based on the resource endowments and economic and social conditions of different spatial scales, and based on the layout of national land space functions and development plans (Huang and He, 2017). Differentiated policies should be developed at different spatial scales to achieve zero emissions and absorption after superposition, ultimately achieving the 2060 carbon neutrality target for the entire national territory (Huang, 2020; Zhang et al., 2021).

In the context of rivers, watersheds form independent ecosystems and economic systems with unique characteristics (Ehrlich and Holdren, 1972). The carbon sources and sinks within the watershed exist in the ecological and economic space of the watershed, forming a unique spatial pattern of carbon neutrality (Huang and He, 2017). For the watershed, the carbon neutrality target does not mean synchronous and zero emissions across the entire region. Instead, it requires a reasonable carbon neutrality spatial organization arrangement based on the spatial distribution of carbon sources and sinks, achieving localized differentiation and overall optimal carbon neutrality. The Yangtze River Basin is an important economic development zone and ecological resource area in China (Huang et al., 2018). Considering the economic development level, carbon emissions level, and carbon sink level of cities in the Yangtze River Basin, there is an obvious spatial overlap of emissions and carbon sinks among cities (Li et al., 2017). The 132 municipal districts in the region exhibit eight types of cities, including high carbon sinks and low emissions in developed areas and low carbon sinks and low emissions in underdeveloped areas (Fu et al., 2008). Some cities can achieve carbon neutrality through their efforts while neighboring cities can form carbon neutrality circles. There are also many cities that need to arrange their carbon neutrality spatially at the whole basin scale, such as the carbon-rich cities in Yunnan Province upstream and the rigid emission cities in Jiangsu Province downstream, which can form cross-regional carbon neutrality complementarity. Focusing on the classification of carbon-neutral cities in the entire watershed and implementing carbon neutrality spatial organization among cities, not only helps to promote the carbon neutrality process in the Yangtze River Basin but also has important exploration and demonstration significance for driving and guiding carbon neutrality practices in other watersheds and even the entire national territory.

## 2. Spatial Characteristics of Urban Carbon Neutrality in the Yangtze River Basin

### 2.1. Spatial Background of Urban Carbon Neutrality in the Yangtze River Basin

Basins are the cradle of human civilization. Under the connection of rivers, basins form independent ecosystems and economic systems with unique characteristics. Within China’s territorial space, the basins of the Yangtze River, Yellow River, and other rivers support the ecological and economic maps of the entire country with specific spatial characteristics of carbon sources and sinks (Tan and Huang, 2008). The Yangtze River Economic Belt is an important economic development zone and ecological resource area in China (Huang et al., 2018). Cities with highly concentrated populations and industries along the river are the main areas of carbon sources (Ehrlich and Holdren, 1971). At the same time, the vast forest resources on both sides of the river are also important carbon sinks for the basin and even the country. Based on the development of the Yangtze River Basin, the Yangtze River Economic Belt has gathered over 40% of the population and economic output on 21% of China’s land area (Huang et al., 2018). It is the most powerful and strategically important development space in China, as well as an important ecological space for maintaining the sustainable development of China’s resource environment. The 132 municipal districts in the Yangtze River Basin have a GDP of 30.5 trillion CNY, accounting for 44.5% of the national GDP (Zhang and Pan, 2020), the locations and zoing are shown in Figure 1.

**Figure 1.**
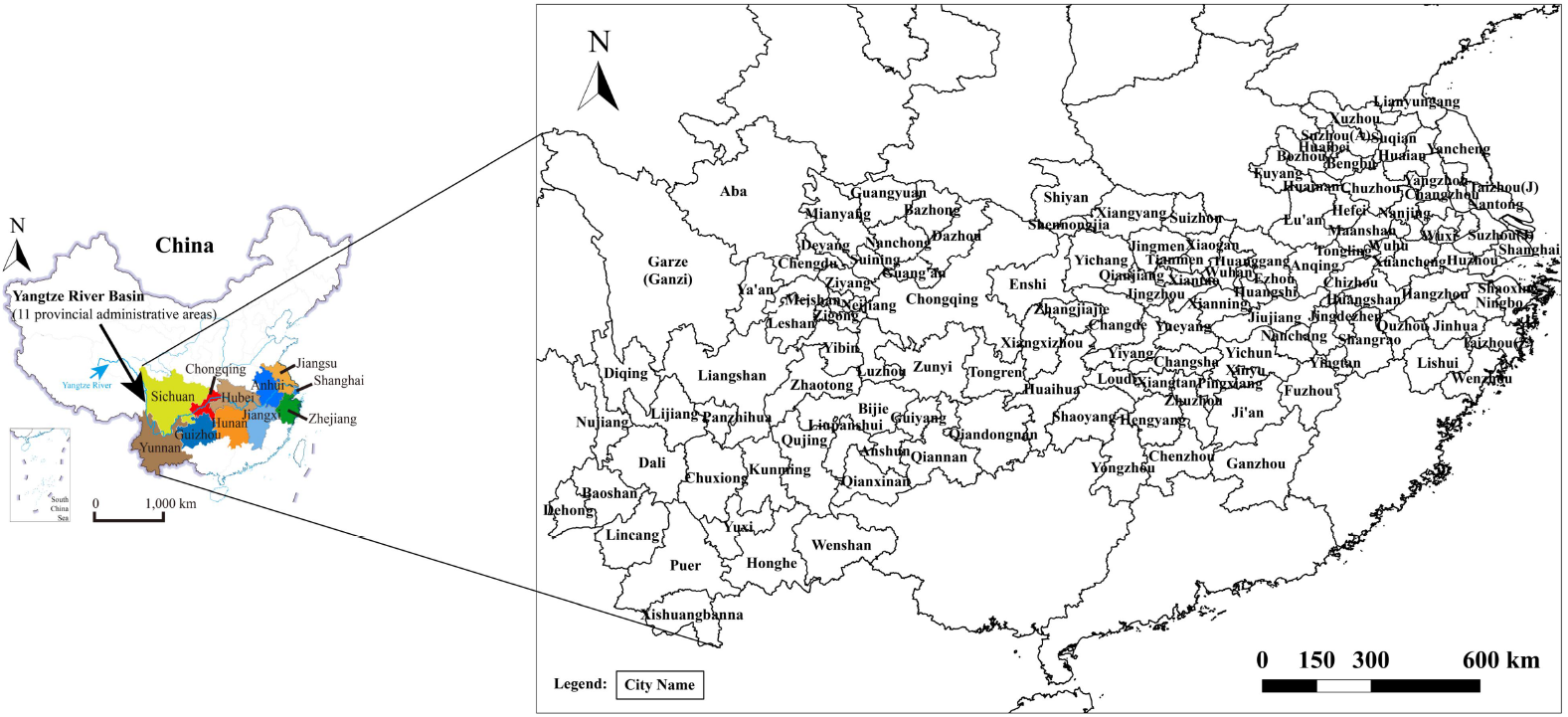
Locations ang zoning of 132 municipal districts in the Yangtze River Basin

The total CO_2_ emissions are 3.886 billion tons, accounting for 32.1% of the total urban emissions in the country. The annual carbon sink absorption is 1.68 million tons, accounting for 33.3% of the total carbon sink absorption in the country. The forest coverage area is 90.47 million hectares, accounting for 41% of the total forest area in the country. Cities in the Yangtze River Basin have achieved 45% of the national GDP with 32% of the national emissions while maintaining a forest cover of 41% and a carbon sink absorption of 33.3% (Huang et al., 2016). Detailed information of the study area is shown in Table 1.

**Table 1.** Statistics of Regional GDP, CO_2_ Emissions and Carbon Sink.

| Region | GRP | CO<sub>2</sub> emissions | Carbon dioxide absorption | Carbon sink emission ratio |
| --- | --- | --- | --- | --- |
|  | billion CNY | billion tons | billion tons |  |
| Yangtze River total | 305200.2 | 38.86 | 3.4825 | 8.96% |
| Yangtze-River proportion | 44.52% | 32.10% | 32.23% |  |
| Shanghai | 25123.45 | 2.77 | 0.0129 | 0.47% |
| Jiangsu | 70116.38 | 7.76 | 0.1147 | 1.48% |
| Zhejiang | 42886.49 | 5.09 | 0.1997 | 3.92% |
| Anhui | 22005.63 | 4.95 | 0.1520 | 3.07% |
| Jiangxi | 16723.78 | 2.45 | 0.3048 | 12.47% |
| Hubei | 29550.19 | 3.26 | 0.2793 | 8.58% |
| Hunan | 28902.21 | 2.86 | 0.3600 | 12.58% |
| Chongqing | 15717.27 | 2.04 | 0.1207 | 5.92% |
| Sichuan | 30053.1 | 3.68 | 0.7232 | 19.66% |
| Guizhou | 10502.56 | 2.31 | 0.2566 | 11.09% |
| Yunan | 13619.17 | 1.70 | 0.9585 | 56.45% |

The Yangtze River Basin is distributed from east to west with various topographies such as coastlines, plains, hills, mountains, and plateaus. The adjacent cities are connected by natural geographical features such as rivers, mountains, shared forests, lakes, wetlands, wildlife migration, and weather phenomena. Against the grand natural geographical background of the Yangtze River Basin, the resource endowments, population size, economic output, level of prosperity, technological level, cultural education, and other factors among the cities in the basin jointly constitute the spatial distribution background that affects the CO_2_ emissions, forest carbon sinks, and carbon neutrality of the cities in the Yangtze River Basin (Li, 2020).

### 2.2. Spatial distribution of carbon neutrality factors in cities in the Yangtze River Basin

From the perspective of the carbon emissions and percentiles of cities in the Yangtze River Basin (Figure 2), the low-emission areas are mainly concentrated in the middle and upper reaches of the Yangtze River, while the majority of cities in the lower reaches have high emissions. The high-emission areas in the upper reaches include Yuxi, Kunming, and Qujing in Yunnan Province, Zunyi, Guiyang, Liupanshui in Guizhou Province, Chongqing City, Guang’an and Dazhou in Sichuan Province, which are generally located near the middle reaches. The large basins to the west are all in low-emission areas. Except for Pingxiang, Ji’an, Fuzhou, Zhuzhou in Hunan Province, Huangshan and Jingdezhen in Anhui Province, Lishui in Zhejiang Province, and Suqian in Jiangsu Province, most cities in the middle and lower reaches of the Yangtze River are high-emission areas. The CO_2_ emissions in the entire basin exhibit a west-low and east-high trend. From the quartile chart (Figure 3), Sichuan Province and a few cities in Yunnan and Guizhou provinces near the boundary of the middle and upper reaches are prominent high-emission areas, while the majority of areas in Jiangsu Province along the coast and some parts of Zhejiang Province are prominent high-emission areas in the middle and lower reaches of the Yangtze River.

**Figure 2.**
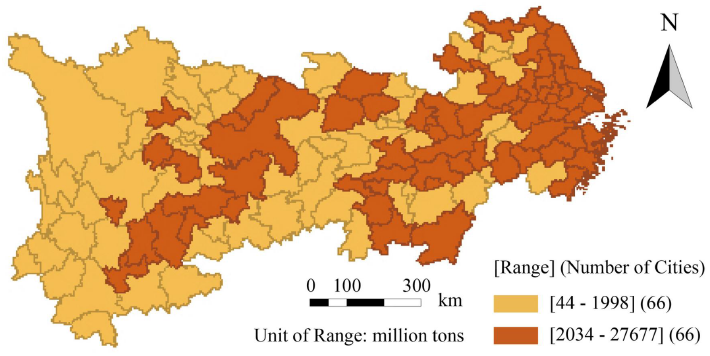
Binary map of carbon dioxide

**Figure 3.**
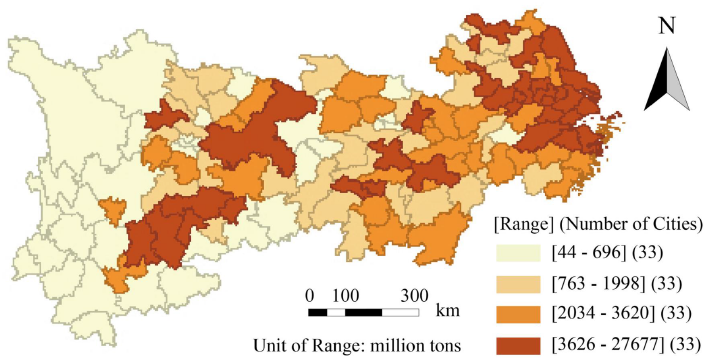
Quartile map of carbon dioxide

From the perspective of the forest coverage and percentiles of cities in the Yangtze River Basin (Figure 4), except for Qujing and Zhaotong in Yunnan Province, Xiangtan and Hengyang in Hunan Province, Jingdezhen and Nanchang in Jiangxi Province, Hangzhou and Ningbo in Zhejiang Province, all of the aforementioned provinces, as well as Guizhou Province, are carbon sinks with relatively high forest coverage. Except for most cities in Hubei Province and Le Shan, Ya’an, Mianyang, Ba Zhong, and Guangyuan in Sichuan Province, all of Sichuan, Shanghai, Chongqing, Anhui, and Jiangsu are regions with low forest coverage. The forest carbon sink in the entire basin exhibits an obvious north-south gradient trend. From the quartile chart (Figure 5), it can be seen that Yunnan Province in the southern part of the upstream, southern Jiangxi Province in the middle reaches, and southern Hunan Province are prominent areas with high forest carbon sinks, while most of Jiangsu Province in the east and some parts of Anhui Province are prominent areas with low forest carbon sinks.

**Figure 4.**
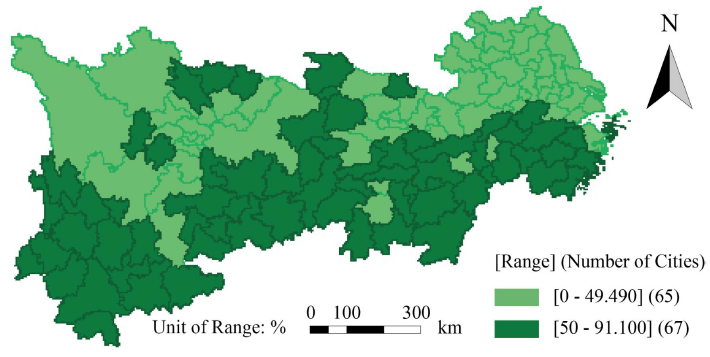
Binary map of urban forest coverage rate

**Figure 5.**
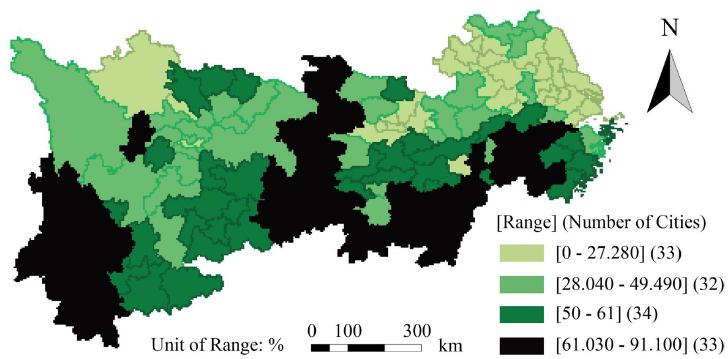
Quarter map of urban forest coverage rate

The quartile chart shows that the distribution of carbon sources in the Yangtze River Basin exhibits a pattern of higher emission in the east and lower emission in the west, while the forest coverage distribution presents a pattern of higher coverage in the south and lower coverage in the north. The emission-to-sink ratio exhibits a pattern of alternating high and low values. The middle-lower reaches south of the Yangtze River have high emission and high carbon sink characteristics, while the middle-lower reaches north of the river have high emission and low carbon sink characteristics. The middle-upper reaches north of the river have low emission and low carbon sink characteristics, and the middle-upper reaches south of the river have low emission and high carbon sink characteristics. The uppermost and lowermost regions show the most significant differences in carbon source emission and forest carbon sinks, making them the areas with the most significant differences between carbon source and carbon sink (Carbon Sink).

To describe the contrast between carbon dioxide emissions and forest carbon sinks in the Yangtze River Basin and to display the level of carbon neutrality pressure, the emission-to-sink ratio is constructed as the ratio of emissions to forest coverage, reflecting the level of carbon neutrality pressure. Regions with high emission-to-sink ratios are emission areas, while those with low ratios are carbon sink areas. From the quartile chart of emission-to-sink ratios of cities in the Yangtze River Basin (Figure 6), the ratio exhibits an obvious low-high alternating distribution along the basin. Regions near the middle reach in the upstream and the transition area between the middle and lower reaches exhibit higher emission-to-sink ratios, as do coastal areas. These regions are interspersed with areas with lower emission-to-sink ratios, demonstrating a pattern of alternating carbon source and carbon sink areas along the basin. From the quartile chart (Figure 7), it can be seen that the uppermost region has the lowest emission-to-sink ratio, while the lowermost region has the highest ratio.

**Figure 6.**
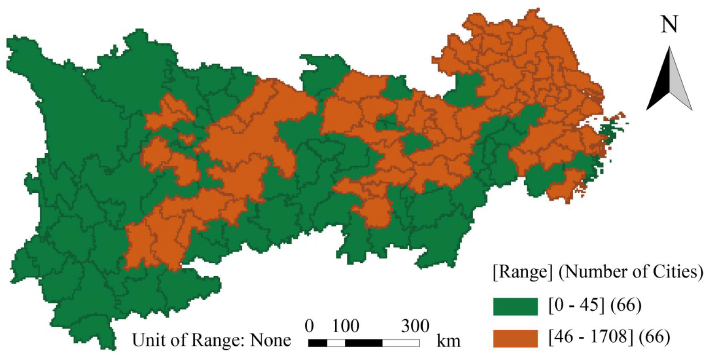
Binary map of CO_2_ and CS ratio

**Figure 7.**
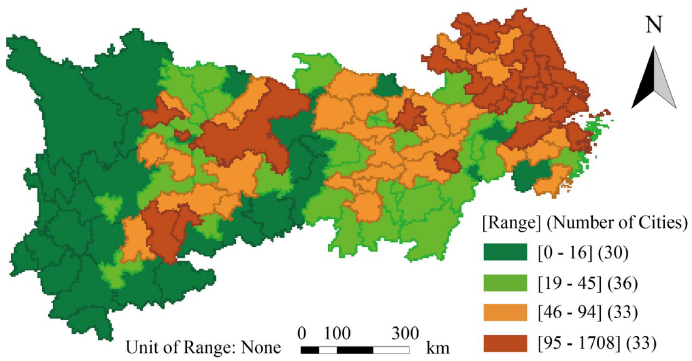
Quarter map of CO_2_ and CS ratio

From the quartile chart of the Gross Domestic Product (GDP) of cities in the Yangtze River Basin (Figure 8), it can be observed that the economic development in the region exhibits an obvious pattern of higher GDP in the east and lower GDP in the west. This pattern is similar to the spatial distribution of carbon dioxide emissions in the main sections of the basin. However, there is also a clear misalignment between high-emission underdeveloped cities and low-emission developed cities. From the quartile chart (Figure 9), it can be seen that the Yunnan province, most parts of Guizhou province, and most parts of Sichuan province in the middle-upper reaches are underdeveloped regions with low emissions. In contrast, most parts of Jiangsu province in the east and a small portion of northern Zhejiang province are developed regions with high emissions. Other regions exhibit a mixed pattern of emissions and development levels. Overall, there is a gradual increase in emissions and development levels from west to east along the Yangtze River Basin.

**Figure 8.**
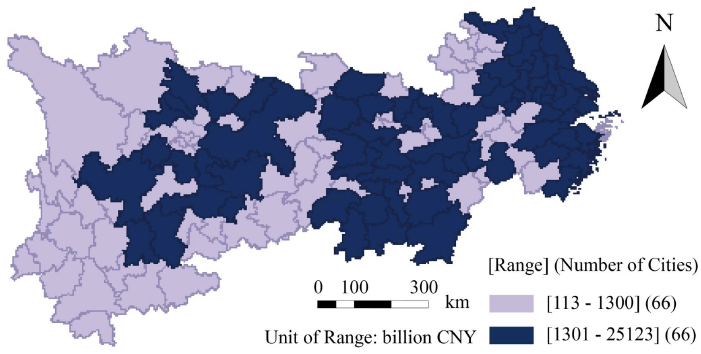
Binary map of GDP

**Figure 9.**
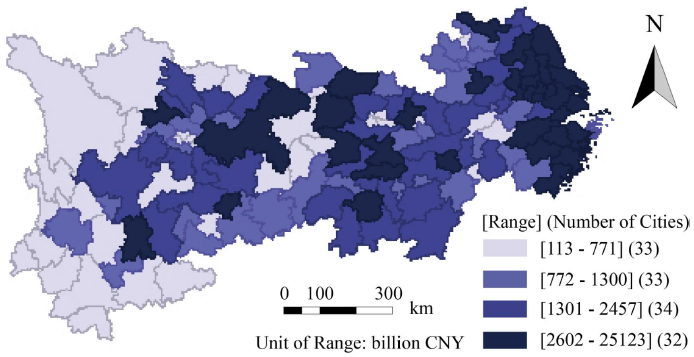
Quarter map of GDP

By combining the quartile charts of carbon dioxide emissions, forest coverage, emission-to-sink ratio, and Gross Domestic Product (GDP) of cities in the Yangtze River Basin (Figures 3, 5, 7, and 9), it can be observed that high-emission developed areas in the basin mainly concentrate in the downstream coastal regions of Jiangsu and Zhejiang provinces, as well as in the western Chengdu-Chongqing region. These areas have advantages in high-carbon industries and production capacity but are not high-carbon sink areas. Low-emission underdeveloped areas in the basin mainly concentrate in some cities of Yunnan province and Sichuan province in the upper reaches, where Yunnan province has higher forest coverage and is a low-emission underdeveloped high-carbon sink area. The Yangtze River Basin is an area with relatively high economic and social development in China, and there are no obvious high-emission underdeveloped areas within the basin. However, despite the relatively high economic and social development in the Yangtze River Basin, there is no low-emission high-carbon sink developed area that has formed a demonstrative influence within the basin.

### 2.3. Spatial Relationship of urban carbon neutrality factors in the Yangtze River Basin

The spatial Gini coefficient can reflect the spatial average degree of distribution of urban emissions, forest carbon sink levels, and emission-to-sink ratios within the basin. The formula is as follows:

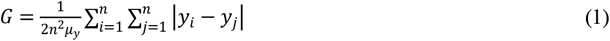

*G* is the Gini coefficient, *y*_*i*_ is the value of the ith city, | *y*_*i*_ − *y*_*i*_ | represents the absolute value of the difference between any two cities, *n* is the number of cities, and *μ* is the basin average value. The Gini coefficient is obtained by summing the differences between cities and dividing by the average of the square of the index for all cities. The Gini coefficient ranges from [0, 1], with values closer to 1 indicating a greater concentration of variables in geography and greater inequality in distribution, while values closer to zero indicate a more even distribution. Spatial Gini coefficient analysis of carbon dioxide emissions, forest coverage, and emission-to-sink ratios in cities in the Yangtze River Basin shows that the Gini coefficient of forest coverage in the basin is 0.2288, indicating a relatively even distribution, while the Gini coefficient of carbon dioxide emissions is 0.5193, indicating a significant level of imbalance, and the Gini coefficient of emission-to-sink ratios is 0.6336, indicating the most significant spatial imbalance (Table 2). Thus, it can be seen that forest resources, which are mainly distributed naturally, are in a relatively balanced distribution, while carbon dioxide emissions resulting from human factors exhibit a high degree of spatial concentration. When considering both emission and forest carbon sink factors, this spatial imbalance is reinforced, resulting in the highest Gini coefficient of emission-to-sink ratios. Carbon neutrality in cities in the Yangtze River Basin is in a significantly spatially imbalanced state, and the uneven distribution of emissions is the main aspect of the spatial imbalance in carbon neutrality in the basin (Yu et al., 2012).

**Table 2.** Relationship table of major indicators of carbon neutrality.

|  | Carbon source emissions | Forest carbon sinks | Emissions to carbon sink<br>ratio | GRP |
| --- | --- | --- | --- | --- |
| Gini coefficient | 0.5193 | 0.2288 | 0.6336 | 0.5411 |
| Global Moran's index | 0.0889 | 0.0688 | 0.0865 | 0.0520 |

Spatial autocorrelation research assumes that things distributed in space are interrelated and that the interaction between nearby things is greater than that between distant ones (Li, 2019). Variables such as emissions and forest coverage rates in a city are affected by surrounding cities and, in turn, affect neighboring cities. Moran’s I can be used to measure the accumulated degree of this influence across the entire basin.

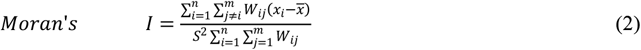

In the formula, *n* represents the number of cities, while *x*_*i*_ refers to the emission, forest carbon sink, and emission-to-sink ratio values of a specific city. 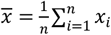 represents Variable mean. 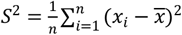 represents the variance of the variable, *m* is the number of cities in the weight matrix, and *Wij* stands for Spatial Weight Matrix. The spatial weight matrix is an abstraction of spatial relationships and a quantification of such relationships. Moran’s I ranges from -1 to 1, and the closer it is to 1, the higher the degree of spatial autocorrelation. The closer it is to 0, the lower the degree of spatial autocorrelation. The closer it is to -1, the higher the degree of negative spatial autocorrelation. In this study, a spatial adjacency weight matrix was used to conduct a Moran’s test on the basin-wide global spatial autocorrelation of carbon emissions, forest carbon sinks, and emission-to-sink ratios in cities in the Yangtze River Basin. The Moran’s index for forest coverage in the basin was 0.0688, the Moran’s index for carbon dioxide emissions was 0.0899, and the Moran’s index for emission-to-sink ratios was 0.0866, all of which were relatively low (Table 2). Therefore, it can be seen that both forest resources, which mainly have a natural distribution, and carbon dioxide emissions resulting from human factors do not exhibit obvious spatial autocorrelation effects from the overall perspective of the Yangtze River Basin. The emission-to-sink ratio, which is closely related to carbon neutrality, also exhibits spatially random distribution characteristics. Therefore, from the perspective of the entire basin, there is relative independence between carbon sources and forest carbon sinks in the Yangtze River Basin.

## 3. Carbon-neutral spatial organization in the Yangtze River Basin

### 3.1. Classification of carbon neutrality types in cities in the Yangtze River Basin

In the process of dealing with climate change, economic development, energy conservation, and emissions reduction have always been interrelated. Carbon neutrality has added the dimension of forest carbon sinks, and different regional characteristics have emerged in the national territory under the joint influence of economic development level, emission level, and carbon sink level. These regions are the basic objects of carbon neutrality spatial organization. As long as there are differences in economic activities and carbon sink distribution among different regions, there will be regional differences in carbon neutrality, which requires the implementation of differentiated regional carbon neutrality strategies based on spatial differences in carbon neutrality. Based on the comprehensive levels of economic development, carbon dioxide emissions, and forest coverage in cities in the Yangtze River Basin, eight types of urban areas can be classified, including high-emission and high-carbon-sink developed areas, high-emission and high-carbon-sink underdeveloped areas, high-emission and low-carbon-sink developed areas, high-emission, and low-carbon-sink underdeveloped areas, low-emission, and high-carbon-sink developed areas, low-emission and high-carbon-sink underdeveloped areas, low-emission and low-carbon-sink developed areas, and low-emission and low-carbon-sink underdeveloped areas (Table 3). These spatial characteristic types outline the spatial layout characteristics of urban carbon neutrality within the basin scope.

**Table 3.**
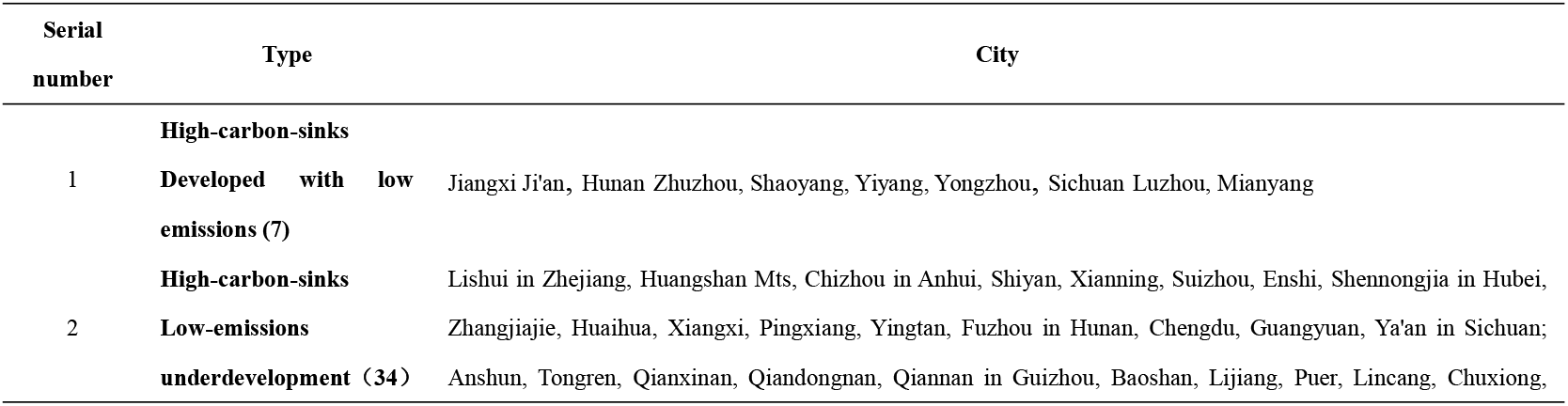
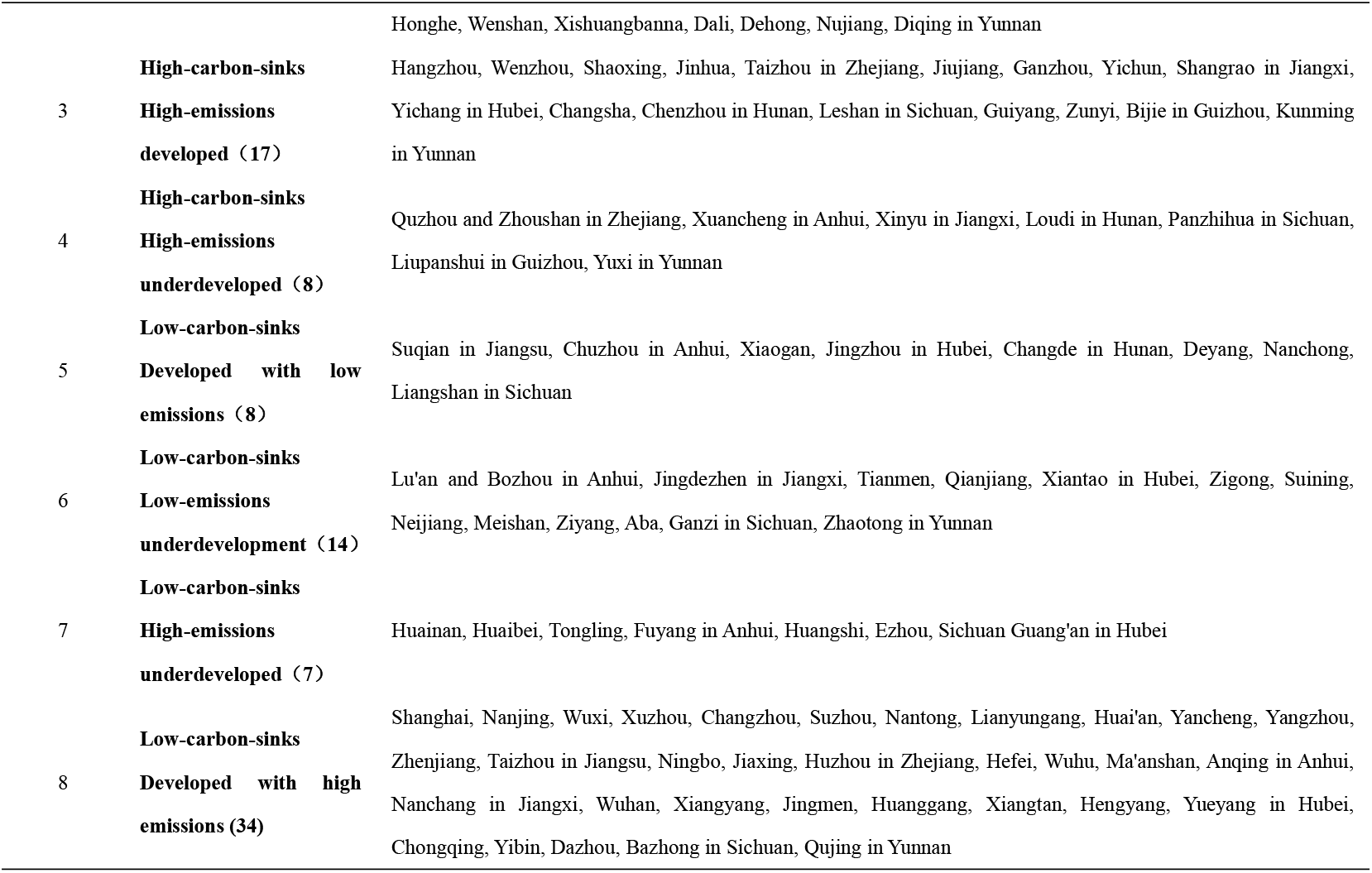
Classification of carbon neutralization types.

In the cities of the Yangtze River Basin, the majority of carbon neutrality types are low-carbon-sink and high-emission developed cities and high-carbon-sink and low-emission undeveloped cities, with 34 cities in each type, accounting for 51.1% of all cities in the basin, while the remaining six types accounting for 48.9%. The main contradiction in the spatial organization of carbon neutrality in the Yangtze River Basin lies in balancing the low-carbon-sink and high-emission developed cities with the high-carbon-sink and low-emission undeveloped cities. The 34 low-carbon-sink and high-emission developed cities in the Yangtze River Basin include the two municipalities of Shanghai and Chongqing, as well as Jiangsu Province in the coastal developed area and Hubei Province in the middle reaches of the Yangtze River. All 12 cities in Jiangsu Province except Suqian belong to the low-carbon-sink and high-emission developed city type, and seven cities in Hubei Province belong to this type, mainly distributed on the north side of the basin, serving as the primary carbon emissions sources within the basin. The 34 high-carbon-sink and low-emission undeveloped cities in the Yangtze River Basin are mainly concentrated in Yunnan Province in the upper reaches with 12 cities, some cities in Guizhou Province, and some cities in Hunan Province in the middle reaches of the Yangtze River. The carbon neutrality of both the low-carbon-sink and high-emission developed cities and the high-carbon-sink and low-emission undeveloped cities within the entire basin consist of two parts. The first part involves cross-spatial emissions and carbon sink neutrality between the developed coastal emission region and the underdeveloped carbon sink region in the upper reaches of Yunnan and Guizhou. The second part is the carbon sink emission neutrality formed by the carbon sinks in Hunan, the emissions in Hubei, and the emissions and carbon sinks in Chongqing in the middle reaches of the Yangtze River. The carbon neutrality of these two parts determines the spatial main scenario of carbon neutrality within the Yangtze River Basin.

### 3.2. Leading by example to promote carbon neutrality in territorial space

Carbon neutrality within the scope of territorial space is faced with more complex spatial relationships (Yuan and Sun, 2020). From the perspective of emissions, highly-emitting developed areas generally concentrate heavy chemical industries that have both emission rigidity and demand rigidity. They are regions with high carbon productivity and large emissions. Highly emitting underdeveloped areas are mostly regions with relatively low levels of economic and social development but with high carbon energy and resource advantages and strong desires for improving people’s livelihoods. Low-emission developed areas have a low-carbon industrial structure and low-carbon energy structure, with higher levels of economic and social development and carbon productivity, and they are regions of low-carbon green development (Qing and Jin, 2917). Most low-emission underdeveloped areas are backward regions with lagging economic and social development, including some high-carbon sink areas with better natural environments and low-carbon sink areas with harsh natural conditions. From the perspective of carbon sinks, high-carbon sink areas include highly-developed high-carbon sink areas and highly-underdeveloped high-carbon sink areas. Highly developed high-carbon sink areas are regions with more carbon sink resources, higher levels of economic and social development, and protected and restored ecological environments. Most highly-underdeveloped high-carbon sink areas have relatively low levels of economic and social development and abundant animal and plant resources, and their ecological environment is well protected. Most of these areas find it difficult to achieve carbon neutrality through their efforts and require the effective spatial organization to achieve global-scale carbon neutrality.

The Yangtze River Economic Belt is China’s most important economic development zone and also a crucial ecological resource repository. The demonstration of achieving carbon neutrality in the Yangtze River Economic Belt and the establishment of a carbon-neutral experimental demonstration zone in the Yangtze River Basin is essential for exploring the spatial organization of carbon neutrality and are of great significance to achieving China’s goal of carbon neutrality by 2060.

Firstly, the “ecological priority” development of the Yangtze River Economic Belt has received great attention from the country’s leadership. General Secretary Xi Jinping has repeatedly emphasized the need to prioritize ecology and green development in the development of the Yangtze River Economic Belt and to focus on protecting the environment while avoiding excessive exploitation. This provides clear guidance for the implementation of carbon neutrality pilot projects in the Yangtze River Basin.

Secondly, the “ecological priority, regional interaction, and intensive development” development concept established in the “Development Plan Outline of the Yangtze River Economic Belt” issued in 2016 sets development goals and provides a framework for the development of carbon neutrality demonstration zones in the Yangtze River Basin.

Thirdly, the Yangtze River Basin is China’s most important economic development belt and ecological resource repository. The region’s GDP and forest coverage both accounts for over 40% of the national total, and its carbon dioxide emissions represent over 30% of the country’s total. Achieving carbon neutrality in the Yangtze River Basin would make a significant contribution to mitigating climate change in China.

Fourthly, the Yangtze River Basin spans the eastern, central, and western parts of China and contains diverse types of carbon-neutral cities. It is home to influential cities such as Shanghai, Chongqing, Wuhan, Chengdu, and Hangzhou and can provide a wealth of guiding experience for carbon neutrality efforts throughout China. Additionally, the provinces and regions of Zhejiang and Guizhou within the Yangtze River Basin are pioneers in ecological compensation and ecological civilization construction and have valuable experience and foundations for further piloting and exploration.

### 3.3. Recommendations for promoting carbon neutrality in cities in the Yangtze River Basin

The developed cities with high emissions and low carbon sinks in the Yangtze River Basin are mainly concentrated on the north side of the lower reaches of the Yangtze River, including the two municipalities of Shanghai and Chongqing, as well as some cities in Jiangsu and Hubei. These cities are constrained by rigid emissions and face difficulties achieving carbon neutrality on their own, making them the main source of carbon emissions in the basin. The underdeveloped cities with high carbon sinks and low emissions are mainly located in Yunnan Province and some areas of Guizhou, which have obvious advantages in carbon sink resources and are the main carbon sink cities in the basin. These two types of cities account for more than half of the total number of cities in the Yangtze River Basin. Those which are close to each other can form a carbon-neutral city cluster, while those which are far apart can form a carbon-neutral alliance and implement corresponding carbon-neutrality strategies. The developed cities with high carbon sinks and low emissions among the other six types of cities in the Yangtze River Basin are the ideal type of carbon-neutral cities. However, there are few such cities in the basin, and no important city has reached the stage of being a developed city with high carbon sinks and low emissions. This suggests that there is still a lot of room for improvement in carbon neutrality in the basin, and the task is still daunting. Some cities have economic and technological capabilities to transform into developed cities with high carbon sinks and low emissions through technological upgrades and energy conservation and emission reduction measures, while others need to change their traditional development ideas and firmly restrain the trend of developing with inefficient energy use and lower carbon productivity. From the perspective of environmental protection and increasing carbon sinks through afforestation, future development goals, and tasks should be planned. Some cities currently have low levels of carbon sinks but have geographical conditions and economic foundations to vigorously develop forestry and transform into developed cities with high carbon sinks and low emissions. Effective measures are needed to promote the development of forestry in these cities and accelerate their transformation into developed cities with high carbon sinks and low emissions. The underdeveloped cities with low carbon sinks and high emissions, as well as those with low carbon sinks and low emissions, are mainly located in the middle and upper reaches of the Yangtze River. The primary task for these cities is to change their views on economic growth achievements and incorporate afforestation and ecological environmental protection as important tasks of economic and social development while steadily improving people’s living standards. These cities should focus on increasing carbon sinks and taking on more carbon neutrality tasks.

The carbon neutrality of the Yangtze River Basin should first rely on the efforts of cities within the basin to improve their own carbon neutrality levels, transform their development concepts, and establish a development concept that regards green mountains and clear waters as golden and silver mountains. They should also change their single-minded pursuit of economic growth and adopt a stance that focuses on improving industrial structures, energy structures, and technological levels, reducing emissions, and vigorously developing forestry to increase carbon sequestration. This will serve as a driving force for their own carbon neutrality and help to improve the carbon neutrality level of the entire basin.

Secondly, cities with low emissions and high carbon sequestration should be established as targets within the basin, serving as demonstrative cities about carbon neutrality. Such cities will contribute to carbon sequestration in the surrounding areas, while demonstrating the effectiveness of the approach and encouraging other cities within the basin to emulate their practices, thus raising the overall carbon neutrality level of the entire basin.

Thirdly, it is essential to strengthen the provision of livelihood guarantees and income improvement in underdeveloped, high-carbon sequestration areas, fortify social support for environmental protection and carbon sequestration resources, firmly suppress the increase of high-emission and low-carbon sequestration cities, and maintain the stability of forest carbon sequestration resources throughout the entire basin.

Fourthly, it is important to enhance carbon neutrality cooperation and communication among cities within the basin, reinforce cooperation between them with regard to emission reduction and afforestation, communicate information on time, negotiate and establish goals, and jointly promote carbon neutrality. This will enable the entire basin, as well as individual cities, to work together in advancing carbon neutrality and develop the new territory as a pioneering and demonstrative zone for spatial carbon neutrality in China.

Fifthly, both government and market should play a dual role from an ecological fairness perspective, promoting spatial carbon neutrality in the basin through transfer payments and government procurement, establishing a carbon emissions trading mechanism in the basin, and promoting market-oriented spatial cooperation between carbon sources and carbon sequestration areas within cities. This should be especially focused on carbon neutrality spatial cooperation policies specifically for upstream underdeveloped cities with high carbon sequestration and downstream coastal developed cities with high emissions and low carbon sequestration, thus realizing coordinated spatial carbon neutrality planning under joint government and market action (Chen et al., 2018; Land Hu, 2019).

Lastly, a specialized organization should be established under the auspices of the National Department of Ecology and Environment to coordinate the carbon neutrality process throughout the Yangtze River Basin. The government should play a leading role in coordinating, advancing, supervising, and guiding carbon neutrality practices throughout the basin, and adopt different approaches for different types of cities and regions. Active efforts should also be made to cultivate the carbon neutrality market within the basin, make carbon neutrality an important criterion for high-quality development of the Yangtze River Basin, and work together to realize the spatial carbon neutrality planning for the entire basin, combined with ecological civilization and the “Beautiful China” initiative.

^1^ Zhang, Y., Wang, X., 2021. Study on forest volume-to-biomass modeling and carbon storage dynamics in China. SCIENTIA SINICA Vitae (02), 199-214.

